# Dose old age lead to gluttony? Effects of aging on predation and locomotion in the assassin bug *Amphibolus venator*

**DOI:** 10.1101/2019.12.16.877993

**Authors:** Kentarou Matsumura, Mana Iwaya, Naohisa Nagaya, Ryusuke Fujisawa, Takahisa Miyatake

## Abstract

Animal behaviors are often affected by aging. In many insect species, locomotor activity decreases with aging. Foraging ability may also decrease with aging. However, few studies have investigated the effects of aging on both locomotor activity and foraging ability. In the present study, we tested the aging effect on locomotor activity and foraging ability in the assassin bug *Amphibolus venator*. The present results showed that locomotor activity decreased with age, similar to findings in many other animal species. However, foraging ability increased with age. Namely, the decline in locomotor activity with age did not lead to a decline in foraging ability. The positive relationship between foraging ability and age may be related to the type of predation, sit-and-wait, used by *A. venator* via alterations in investment in reproductive traits with age.

## Introduction

Aging is a progressive natural process in which deterioration of physiological function occurs [1–3]. Many previous studies have reported that behavioral performance often decreases with age in vertebrate and invertebrate animals [4–10]. Among invertebrates, the fruit fly *Drosophila melanogaster* has been used extensively to study age-related behavioral changes [5]. Previous studies have focused on the decline in behavioral traits with age in flies, including duration of flight [7] and locomotor activity [8,9], and behavioral performance may suffer negative effects of aging even in insect species. In insects, negative effects of aging are considered to occur due to damage in appendages, including the legs and the cuticula that make up their structure [10].

This loss in mobility could influence an animal’s ability to acquire resources. Food intake is a vital function that decreases with age. In mammals, aging is also associated with declines in food consumption [11,12]. Increased age has also been shown to be associated with a decrease in foraging efficiency in some invertebrate species [5,13,14]. However, few studies have examined the link between locomotor abilities and foraging efficiency as a result of aging effect in animals [but see 15].

In the present study, we tested the effects of aging on locomotor activity and foraging ability in the assassin bug *Amphibolus venator* (Klug) [Reduviidae]. *A. venator* is a carnivorous insect, and they often eat stored-grain insects, including the red flour beetle *Tribolium castaneum* [16]. We hypothesized that if *A. venator* also shows decreased locomotor activity with age, then foraging ability could decrease with age as well. To test this hypothesis, we investigated locomotor activity and foraging ability using *A. venator* adults of various ages.

## Materials and Methods

### Insect and culture

The population of *A. venator* used for the present study was collected in Urasoe City, Okinawa, Japan, in 2015, and this population has been maintained in the laboratory of Okayama University [19]. Each bug was reared in an incubator maintained at 29°C and 16L:8D (light on at 7:00, light off at 23:00) light conditions with *T. castaneum* as food.

### Locomotor activity

To measure the locomotor activity of *A. venator*, we used a treadmill system, ANTAM [18]. The ANTAM is developed from the omnidirectional treadmill mechanism system in which animal movements can be continuously recorded and compensated for in such a way that the animal is always located on the top of the sphere and experiences a virtual unbounded two-dimensional field [18]. Therefore, this system is able to measure the free walking trajectories of small animals such as insects [19]. The walking speed of *A. venator* is 26.60 ± 15.41 mm / sec (mean ± s.d., n = 133; unpublished), which is within the allowable range of the movement speed of the system used (for example, ANTAM can measure the walking speed of 55.2 ± 34.3 mm/sec (average ± s.d.) in the pill bug, *Armadillidium vulgare* [18]).

Virgin males (*N* = 59) and females (*N* = 74) were randomly collected, and each bug was placed on the ANTAM system. When a bug was moving, we recorded locomotor activity for 10 min. To examine the effects of starvation on movement (i.e., foraging activity), we also measured the locomotor activity of bugs with subjected to a starvation treatment for 7 days (male: *N* = 10, female: *N* = 10) and fed bugs (male: *N* = 10, female: *N* = 10). Measurements were conducted between 10:00 and 18:00 in a room maintained at 25 ± 2°C.

### Predation

We measured the foraging ability of *A. venator* in small- and large-scale containers over 10 days because of the possibility of a difference in the opportunities for foraging between bugs in small- and large-scale containers. In the small-scale container experiment, virgin males (*N* = 19) and females (*N* = 37) were randomly collected, and each bug was placed into a cylindrical container (35 mm in diameter, 10 mm in height). All bugs were not provided food for 7 days before the experiment. Five *T. castaneum* adults were randomly collected from the stock culture and put into each Petri-dish along with, an *A. venator* adult, and we counted the number of beetles in each Petri-dish that was eaten by the predatory bug every 2 days. If beetles died by predation, we replaced them with live beetles; that is, the density of beetles was kept constant during this experiment.

In the large-scale container experiment, virgin males (*N* = 17) and females (*N* = 24) were randomly collected, and each bug was placed into each cylindrical container (149 mm in diameter, 65 mm in height). All bugs were not provided food for 7 days before the experiment. We measured the number of beetles preyed on by *A. venator* using the same methods as in the small-scale experiment. All predation experiments were conducted in the incubator described above.

### Statistical analysis

In the analysis of locomotor activity, we used three typed data from the ANTAM system: (a) total walking distance (travel distance), (b) direct distance from start to goal (straight distance), and (c) straight distance divide by travel distance (straight rate). To analyze the three data types, we used analysis of variance (ANOVA) with age, sex, and the interaction between age and sex as explanatory variables. To test of the effects of the starvation treatment (treatment) on locomotor activity, we used ANOVA with age, treatment, sex, and the interaction between treatment and sex as explanatory variables.

In the analysis of foraging ability, we used a generalized linear model (GLM) with a Poisson distribution. In this analysis, age, sex, and the interaction between age and sex were used as explanatory variables. Analysis of foraging ability was conducted separately for small- and large-scale experiments.

All analyses were conducted by JMP Ver.12.2.0 [20].

## Results

Figure 1 shows the locomotor activity results. ANOVAs showed significantly negative effects of age on travel distance and straight distance (Fig. 1, Table 1). There were no significant effects of sex and the interaction on either trait. In the straight rate results, there were no significant effects for all factors (Table 1).

**Figure 1.**
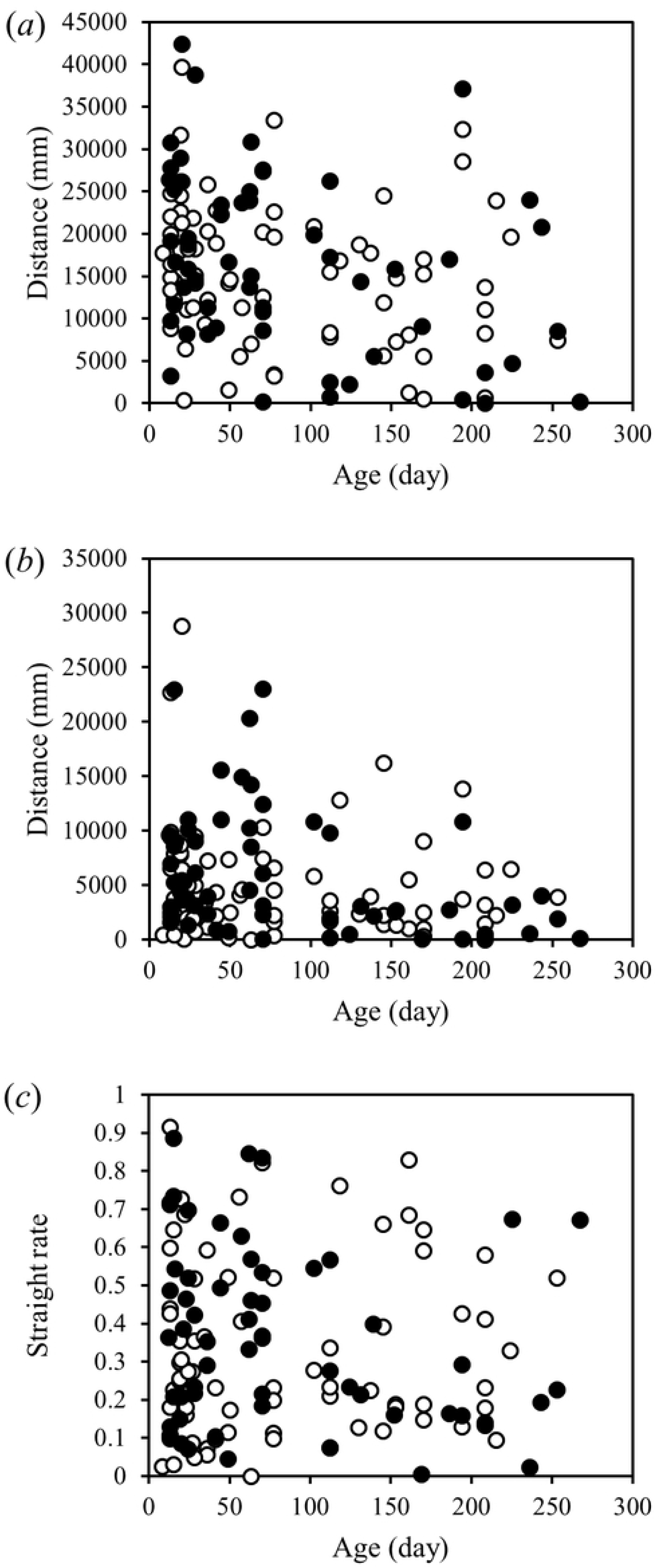
Relationships between walking traits (*a*: travel distance, *b*: straight distance, *c*: straight rate) and age in *A. venator*. Filled and open circles show males and females, respectively.

**Table 1.** Results of ANOVA for walking traits in *A. venator*.

| Trait | Factor | *d.f.* | *F* | *p* |
| --- | --- | --- | --- | --- |
| Travel distance | Sex | 1 | 0.52 | 0.4731 |
|  | Age | 1 | 9.38 | **0.0027** |
|  | Sex*Age | 1 | 1.23 | 0.2696 |
|  | Error | 135 |  |  |
| Straight distance | Sex | 1 | 0.88 | 0.3493 |
|  | Age | 1 | 9.12 | **0.0030** |
|  | Sex*Age | 1 | 1.29 | 0.2586 |
|  | Error | 135 |  |  |
| Straight rate | Sex | 1 | 0.06 | 0.8117 |
|  | Age | 1 | 1.57 | 0.2131 |
|  | Sex*Age | 1 | 1.54 | 0.2169 |
|  | Error | 135 |  |  |

Figure 2 shows the results of locomotor activity including the effect of the starvation treatment. ANOVAs showed no significant effects of treatment on each movement trait (Fig. 2, Table 2). This analysis also showed significant effects of age on travel distance and straight distance (Table 2). There were no significant effects of sex and the interaction between treatment and sex, and all factors did not show effects on the straight rate (Table 2).

**Figure 2.**
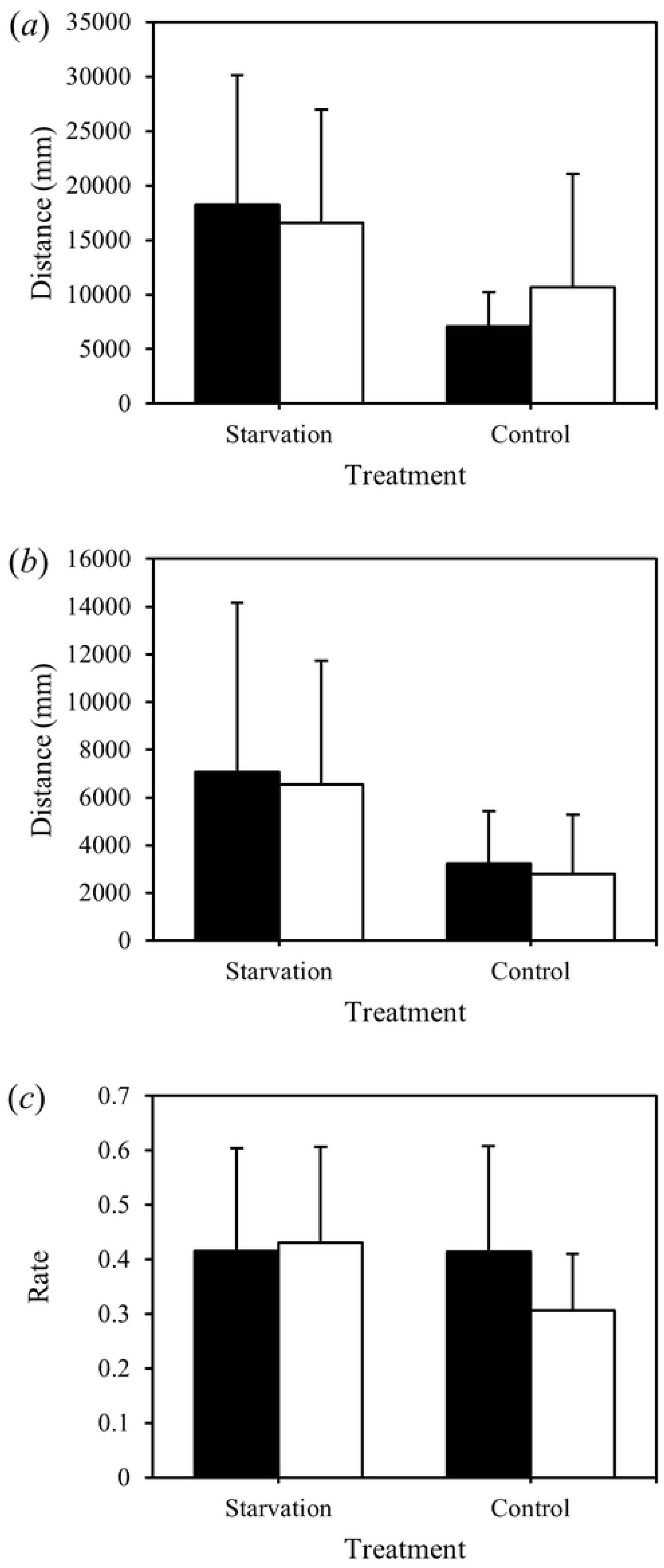
Results of starvation treatment on walking traits (*a*: travel distance, *b*: straight distance, *c*: straight rate) in *A. venator*. Filled and open bars show males and females, respectively. Error bars show the standard error.

**Table 2.** Results of ANOVA for walking traits including effect of starvation treatment in *A. venator*.

| Trait | Factor | *d.f.* | *F* | *p* |
| --- | --- | --- | --- | --- |
| Travel distance | Treatment | 1 | 1.8904 | 0.1779 |
|  | Age | 1 | 8.931 | 0.0051 |
|  | Sex | 1 | 0.4577 | 0.5032 |
|  | Treatment*sex | 1 | 0.2098 | 0.6498 |
|  | Error | 35 |  |  |
| Straight distance | Treatment | 1 | 1.4415 | 0.238 |
|  | Age | 1 | 7.4898 | 0.0097 |
|  | Sex | 1 | 1.4736 | 0.2329 |
|  | Treatment*sex | 1 | 0.1301 | 0.7205 |
|  | Error | 35 |  |  |
| Straight rate | Treatment | 1 | 0.553 | 0.4621 |
|  | Age | 1 | 0.3384 | 0.5645 |
|  | Sex | 1 | 0.8948 | 0.3507 |
|  | Treatment*sex | 1 | 1.3304 | 0.2566 |
|  | Error | 35 |  |  |

Figure 3 shows the results of the predation experiment at the small- and large-scale, respectively. In the small-scale experiment, there was no correlation between predation and age (Fig. 3a, Table 3). Females showed significantly higher foraging ability than males (Table 3). In the females, there was a positive correlation between predation and age (Fig. 3a). There was a significant effect of the interaction between sex and age on predation (Fig. 3a, Table 3). In the large-scale experiment, there was a significantly positive correlation between predation and age (Fig. 3b, Table 3) and no significant effects of sex and the interaction (Fig. 3b, Table 3).

**Figure 3.**
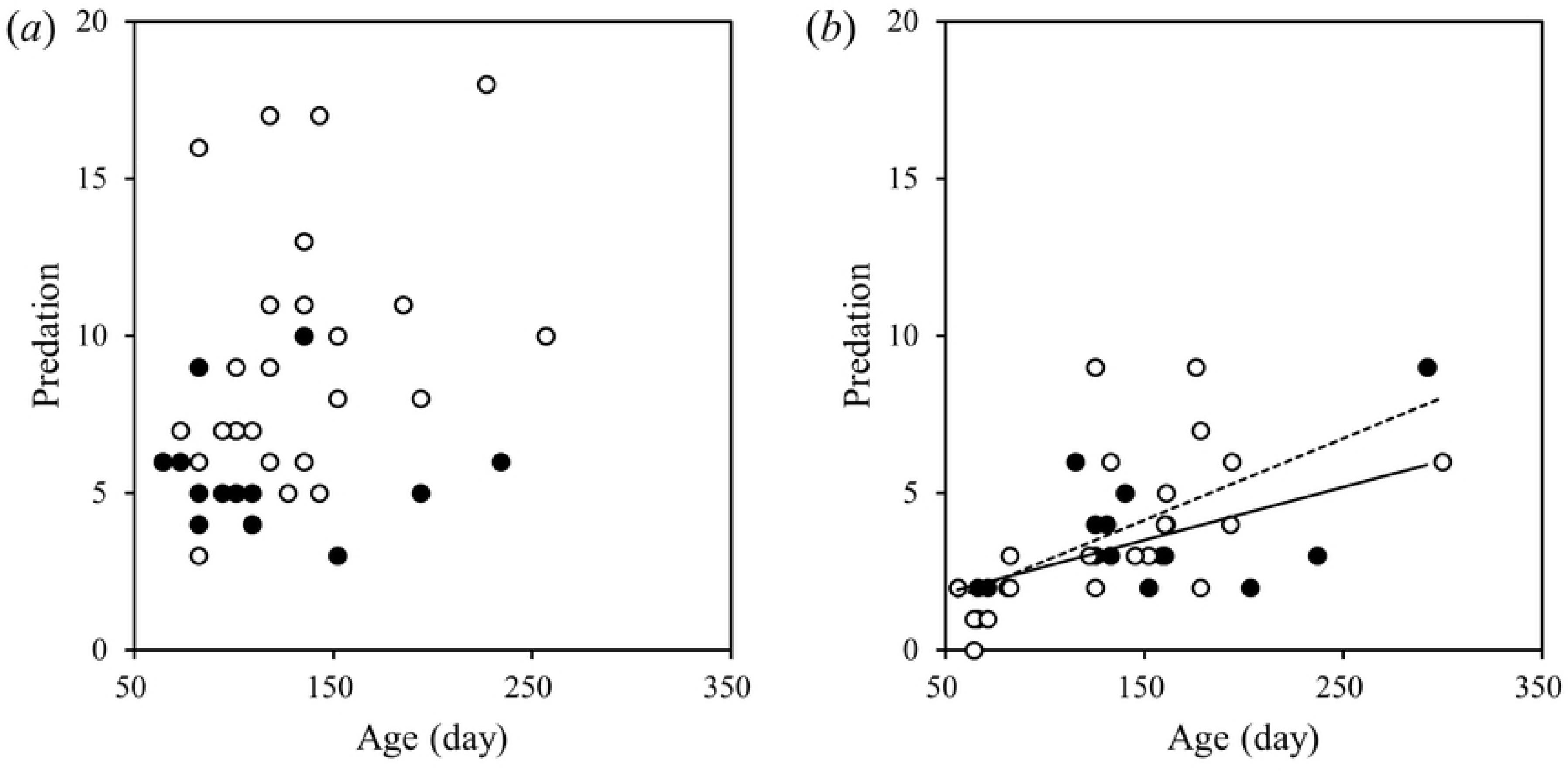
Relationship between age and predation ability in small (*a*) - and large (*b*) -scale experiments, in *A. venator*. Filled (and solid line) and open circles (dashed line) show males and females, respectively.

**Table 3.** Results of GLM for predation in small- and large-scale experiments.

| Scale | Factor | *d.f.* | $\chi^2$ | *p* |
| --- | --- | --- | --- | --- |
| Small | Sex | 1 | 10.61 | 0.0011 |
|  | Age | 1 | 0.68 | 0.4113 |
| | Sex × Age | 1 | 5.37 | 0.0204 |
|  | Error | 52 |  |  |
| Large | Sex | 1 | 0.89 | 0.3452 |
|  | Age | 1 | 13.99 | 0.0002 |
| | Sex × Age | 1 | 0.17 | 0.6822 |
|  | Error | 37 |  |  |

## Discussion

In the present study, locomotor activity was decreased with age, as found in other animal species [5,11–14]. However, foraging ability was not negatively correlated with age. In the small-scale experiment, a positive correlation between predation and age was found only in females. Moreover, in the large-scale experiment, bugs of both sexes showed positive correlations between predation and age. To our knowledge, no study has reported that the older insects are gluttonous.

The decline of locomotor activity with age has often been shown in some animal species [1,21], including in *Drosophila* [5,9,22] and other insect species [10]. In the present study, older bugs showed lower locomotor activity than younger bugs in *A. venator*. Therefore, the present results for *A. venator* are in accordance with those of other previous studies [1,5,8–10,21,22]. In insects, the negative effects of aging are considered to be caused by damage to appendages constructed from cuticula [10]. That is, the present result that locomotor activity decreased with age may be a result of damage to the legs of *A. venator*. Therefore, we need additional studies investigating how damage in appendages relates to age in *A. venator*.

Locomotor activity of *A. venator* did not show an effect in the starvation treatment. A previous study reported that hungry carnivorous bugs, *Deraeocoris lutescens*, showed different walking loci than fed individuals [23]. This difference between previous results and present results may be affected by differences in starvation resistance among the species. The starvation resistance of *A. venator* is relatively high, and we observed that some bugs lived for over 30 days without food (Matsumura personal observation). In the present study, we used bugs that had been held without food for 7 days as starved individuals. This treatment may not be sufficient for *A. venator*. However, there is a large variation in starvation resistance in a population, and some bugs die by from a longer duration of starvation treatment (Matsumura personal observation). We need additional studies to investigate the effects of individual differences in starvation resistance on locomotor activity in *A. venator*.

Some previous studies reported that foraging ability also decreases with age [5,11–14]. However, the present predation experiment showed that the foraging ability of *A. venator* was not negatively associated with age. Therefore, the present results do not agree with those of previous studies. This suggests that foraging ability is not affected by aging in *A. venator*. These differences in results may be caused by the foraging type the predator used in the present study: *A. venator* is a sit-and-wait predator, which is a strategy of waiting until prey approach the predator, and does not actively search for prey [24]. In active searching predators, a decline in locomotor activity is expected to have negative effects on foraging success. On the other hand, in sit-and-wait predators, the foraging ability may not be affected by aging, even when locomotor activity decreases with age. A previous study that used the orb-web spider *Zygiella x-notata*, which is a sit-and-wait predator, showed that the foraging rate was not decreased with age, although the foraging speed was decreased with age [15]. To reveal the association between aging, foraging ability and the type of predation, we need additional studies that investigate foraging ability with age in predators of both the sit-and-wait and active searching types.

Furthermore, the older bugs showed higher foraging ability in the large-scale experiment than the younger bugs. Females also showed a positive correlation between foraging ability and age, even in the small-scale experiment. This result suggests a change in life-history strategy with age in *A. venator*. In many animals, although younger animals invest more in survival than in reproduction, elderly animals invest more in reproduction rather than in survival [25]. Therefore, if *A. venator* also increases investment in reproduction with age, the older individuals may require more resources for gamete production. However, males did not show a correlation between foraging ability and age in the small-scale experiment. This result suggests that males, who may not require more resource than females, did not show a clear association with age because the density of prey was high in the small-scale experiment.

In the present study, there is a possibility that the older bugs were not old enough. In a previous study [17], we investigated the longevity of *A. venator*, and the average longevity was 282.2 days (*s.e.* = 6.78, *n* = 246). In the present study, the number of individuals over 141 days as half the average lifespan was 11/56 individuals in the small-scale and 18/41 individuals in large-scale experiments. We note that although the previous study measured longevity at 25°C, while the present study was conducted at 29°C. That is, the problems associated with age may have been relatively minimal in the present results.

In conclusion, we found that locomotor activity decreased with age in *A. venator*. On the other hand, we showed that foraging ability increased with age. Therefore, these results suggested that the decline of locomotor activity with age was not affected by foraging ability. Moreover, we also suggest that an increase in investment in reproduction with age may lead to a higher foraging ability of older bugs. The present study revealed that aging effects are not decreased for all measures of behavioral performance.

## Acknowledgements

We thank to Mr. Masaya Asakura for reared of *A. venator* in laboratory. This work was supported by a grant from the Japan Society for the Promotion of Science KAKENHI 17H05976 and 18H02510 to TM.

